# Hierarchical Ordination, A unifying framework for drivers of community processes

**DOI:** 10.1101/2024.01.08.574701

**Authors:** R.B. O’Hara, B. van der Veen

**Affiliations:** Department of Mathematical Sciences, Norwegian University of Science and Technology, Trondheim, Norway; Centre of Biodiversity Dynamics, Norwegian University of Science and Technology, Trondheim, Norway; Department of Landscape and Biodiversity, Norwegian Institute of Bioeconomy research, Trondheim, Norway

**Keywords:** Ordination, JSDM, GLLVM

## Abstract

Ordination methods have been used by community ecologists to describe and explore the communities they sample by reducing this variation down to a small number of dimensions. More recently, Joint Species Distribution Models have been developed to model and predict the distributions of several species simultaneously. Contemporary models for the data for both of these problems are essentially the same, called Generalised Linear Latent Variable Models (GLLVMs). Based on this we suggest some avenues of cross-fertilisation between the two areas of research. We also describe some of the extensions to GLLVMs, and from this suggest the development of Hierarchical Ordination, as a way of efficiently modelling communities of species in space.

## Introduction

Community ecology is the study of groups of species. One important set of questions relate to how and why species are distributed relative to each other. For example, do particular species tend to occur together, and can this be explained by similar responses to the environment. These same questions can be asked across different scales, e.g. looking at samples from different Dutch sand dunes (Batterink 1983), or the complete distributions of different species across the globe. Statistical methods have been developed to look at this type of data for both of these scales, but their similarities have not been widely appreciated.

The methods we consider in this article assume that species are observed at a number of sites, so the data is a site by species matrix, with entries being a measure of abundance or incidence, e.g. whether a species was observed at that site, or the number of individuals observed. Historically, these data have been summarised by **ordination methods** (e.g. Gower 1966; Cajo J. F. ter Braak 1985), which look to reduce the variation in data to a small number (usually two) of dimensions. This means that sites and species can be plotted on an **ordination diagram**, which can be inspected visually (Gabriel 1971). More recently, approaches have been developed that are based on modern statistical modelling ideas, leading to the application of Generalised Linear Latent Variable Models in community ecology (Hui et al. 2015; Skrondal and Rabe-Hesketh 2004).

Species distribution models (SDMs) were developed to predict the incidence of species with environmental variables. The perceived importance of interactions between species (Wisz et al. 2013; Kissling et al. 2012) suggested that their distributions should be modelled jointly. This lead to the development of Joint Species Distribution Models (JSDMs: Pollock et al. 2014), which also model the response of each species to the environment, but then incorporate a correlation matrix to allow for covariance between species. It was quickly realised that this matrix becomes unmanageable for many species, as it has so many parameters to estimate. Thus, extensions were developed to model the matrix as the sum of a smaller set of linear effects (e.g. Warton et al. 2015; Ovaskainen, Tikhonov, Norberg, et al. 2017). Because these models are based on a formal probabilistic model (which can be written as a likelihood, and thus fitted to data with flexible statistical methods), they have been extended in a variety of directions (see below).

The differences in the ways that ordination and JSDMs have been used largely revolve around the aims of the data collection and analysis. Ordination has typically been used for small scale field studies that describe the differences between communities they have sampled at different sites or times, and possibly look at how these differences are related to environmental variables (e.g. such as Bray and Curtis 1957). Thus the focus has been on exploration of the relationships between species and communities. In contrast, JSDMs were developed by and for macro-ecologists, wanting to look at the distributions of species over their whole range, with the primary aim of estimating the effects of environmental covariates on the distribution of species, and thus being able to predict their current and future distributions (Clark et al. 2014). The role of the residual correlation was to improve the predictions, rather than be interpretable: problems with interpreting the correlations as interactions between species were pointed out early on (Pollock et al. 2014).

Thus, JSDMs tend to focus on a larger spatial scale, and more emphasis is placed on understanding the effects of covariates. As a practical matter, they are also generally based on binary observations (but see Björk, Hui, et al. 2018 for a counter-example), whereas ordination has been carried out on a wider range of data, and typically at a smaller spatial scale. Despite these differences, the underlying idea - to reduce covariation down to a smaller number of dimensions - is the same. Our aim here is to outline the similarities between the two approaches, and to suggest that each can learn from the other. We then suggest a scheme to flexibly extend these methods without making them too unwieldy.

## The Models

In order to look at the similarities between ordination and JSDMs, we first need to have a common framework. We will thus first develop a simple example of model-based ordination and JSDMs, and use this to explain the more complex models that have been developed.

In the simplest case, the data are a matrix, with rows being sites, and columns being species. The entries are observations of the species, which can take several forms, e.g. counts, percent cover, or (particularly for JSDMs) species’ presence/absence. In addition to this, we can have covariates which are associated with the rows or columns, such as measures of the environment (temperature, habitat etc.) for each site, or traits (size, diet etc.) for each species.

We can create a straightforward model for this data. Each observation of species *j* (*j* = 1, …, *p*) at site *i* (*i* = 1, …, *n*) is denoted *y*_*ij*_. We call the full matrix of observations **Y**. We can model **Y** as an extension of a generalised linear model, with *g*{*E*(*y*_*ij*_)} = *η*_*ij*_, where *g*(·) is a link function (e.g. logit or cloglog for binary observations). On the link scale we model the expected value of each datum as

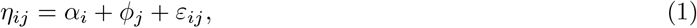

where *α*_*i*_ is the effect for site *i, ϕ*_*j*_ is an effect of species *j*, and where *ε*_*ij*_ is an error term. The error ensures that we incorporate the correlation between species (Pollock et al. 2014):

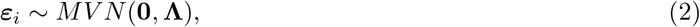

where **Λ** is a *p* ×*p* covariance matrix, with off-diagonal terms being the covariances. The number of parameters in **Λ** increases quadratically with the number of species, which makes the matrix difficult to estimate for large *p*, and the large number of parameters also makes the matrix difficult to interpret. The ordination/JSDM approach to handling this is to write the matrix as the sum of a product of row- and column- effects. Each product defines one of *L latent dimensions*. The model is then

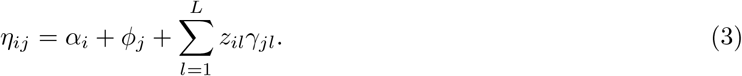

In ordination, *z*_*il*_ is called a site score (i.e. a site its location on the latent variable), and *γ*_*jl*_ is called a species loading. We can interpret *z*_*il*_ as a location along gradient *l*, or informally that there are *L* unobserved covariates, with *γ*_*jl*_ being the regression parameter for the effect of covariate *l* on species *j*. The scale is arbitrary, so it is convenient to scale the site scores so that var(*z*_*il*_) = 1. then the covariance between species one and two becomes 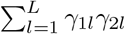.

There are several technical issues that have to be dealt with when fitting this model, for example with two or more latent variables, the site scores are axes that can be rotated, so some constraints are needed so that a unique solution can be found (i.e. to make the model identifiable). However, we will not delve into these details further here.

This is the basic model that has been developed for both JSDMs and ordination (as a GLLVM). As described below, it can be extended in many ways, by including further effects for the sites, species and latent effects, i.e. *α*_*i*_, *ϕ*_*j*_, *z*_*il*_, and *γ*_*jl*_.

### Adding Covariates

The model described above projects the correlation matrix down into *L* dimensions, where *L* is relatively small. But, as written, it does not include covariates. There are several ways to add them: for example, Ovaskainen, Tikhonov, Norberg, et al. (2017) outlined an approach where they incorporate covariates for species and sites respectively, i.e. we could use a model such as:

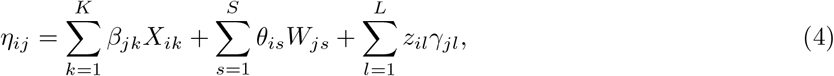

where there are *K* site-level covariates (e.g. climate variables), and *S* species-level covariates (e.g. traits). The model is a sum of several terms, each of which is the product of a species and site term. The differences are whether the terms are known (i.e. covariates) or have to be inferred from the model. So the model incorporates responses of species to environmental conditions, as well as the effects of species’ characteristics on their site response.

The regression coefficient matrices *β* and *θ* can become large if there are many covariates, so some way to reduce this is desirable. This is simply a variable selection problem, for which there are several solutions available, through either selecting which variables are “in” and “out”, or some form of regularisation (e.g. O’Hara and Sillanpää 2009 for some Bayesian methods; or Tredennick et al. 2021 for some non-Bayesian approaches).

Random effects can also be added. This can be interpreted as simply putting a covariance structure on some of the *β*_*jk*_’s or *θ*_*is*_’s. For example, a phylogenetic effect can be added by having a *θ*_*is*_ for each species, but forcing a phylogenetic correlation, i.e. species with a more recent common ancestor tend to have more similar values of *θ*_*is*_.

Classical ordination takes a different approach to including covariates. Instead of the site main effects being modeled, covariates are included through **constrained ordination** (e.g. Cajo J. F. ter Braak 1986), where the site scores of the latent variables are forced to be linear functions of covariates

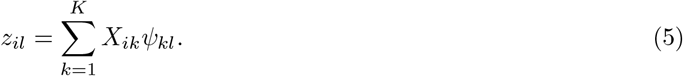

This implies that the latent variables are fully explained by the observed covariates. van der Veen, Hui, Hovstad, and O’Hara (2021) extended this approach to allow a latent variable to be affected by covariates, plus additional (unobserved) effects:

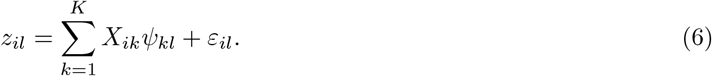

This assumes that the gradient is affected by (or, at least, correlated with) measures of the environment. Again, random effects can also be added, in particular the gradient can be modelled spatially as a continuous field, leading to a type of **spatial factor analysis** (Thorson et al. 2015), where communities that are closer to each other tend to have more similar site effects. In essence, both fixed and random effects can be used to put structure on *z*_*il*_. Another variation on this is to use more than one set of latent variables, with some responding to the environment, and others being unconstrained. For example, Björk, Hui, et al. (2018) modelled host-associated microbiota by including separate ordinations for the host species and the samples (within host species).

The **response** of a species to the gradient can also be modelled as being affected by the environment (e.g. Tikhonov et al. 2017; Perrin et al. 2021):

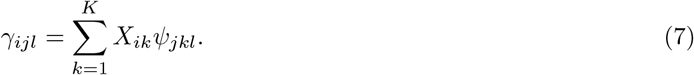

This allows the species effect to depend on the environment, so that the covariance between species *j* and *h* on site *i* is proportional 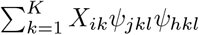, i.e. is a linear function of the covariate. This allows the correlation to change across environments, and potentially change sign.

## Learning From Each Other

Within the development of the GLLVM framework, we can see that JSDMs and model-based ordination are equivalent mathematically. The biological differences stem from the data and the questions being asked. Ordination has typically looked at a set of sites that have been sampled, and asks about the similarities between the communities on these sites. JSDMs, in contrast, try to model and predict the distributions of the species across their whole ranges, i.e. they are interested in the full distributions of the species, and are not interested in characterising the sites themselves. So for ordination, the focus is on the relationship between communities and sites, whereas JSDMs are primarily intended to predict the distribution of each species: the covariance between species is mainly of interest because it changes the predictions for each species (Wilkinson et al. 2021).

The equivalence between ordination and JSDMs suggests that each area should be able to help the other. At the conceptual level, the questions being asked in the ordination world can also be asked by the SDM world, opening up the possibility of modelling the distributions of ecosystems. Most usefully, the methods and theory developed in one area can be transferred across and used in the other.

## What can JSDMs learn from Ordination?

The focus of ordination has usually been on the whole community, rather than looking at individual species. Because of this, the typical summaries are visualisations, i.e. ordination plots, which efficiently condense information from sites and species into figures. In an ordination diagram, sites that are negatively correlated are located far away from each other, and species that are predicted to co-occur will be close together in the figure. These can thus help with understanding co-occurrence patterns, and help to guide further modelling and interpretation (e.g. if several species cluster together). Using ordination plots into the analysis of JSDM should thus help with summarising and interpreting the correlations between species and sites.

The ecological interpretation of ordinations is also more advanced. If the ordination axes are interpreted as ecological gradients, then the site scores represent the predicted locations of sites on the gradient, and the species loadings can (in some models) be the species’ optima. Thus the ordination axes can be interpreted as part of the species’s niche. Ecologically, the development and application of ordination methods has been motivated by the species packing model (Jamil and Ter Braak 2013), where each species has an equal tolerance to the gradient. Adding a quadratic term in the ordination can relax the equal tolerance assumption, so species can be generalists or specialists with respect to the gradient (van der Veen, Hui, Hovstad, Solbu, et al. 2021). On top of this, a constrained ordination can be used to efficiently model the niches of many species together. This will be useful when a large number of species are being considered, e.g. from meta-barcoding data, and particularly when properties of the whole community, rather than of each species, will be important.

## What can Ordination learn from JSDMs?

Ordination can take advantage of the flexibility of the modelling framework, which has been developed more fully in the JSDM world. As described above, Ovaskainen, Tikhonov, Norberg, et al. (2017) developed a framework to incorporate a wide range of effects on both the species and sites, which can potentially incorporate spatial effects (Tikhonov et al. 2020). The statistical modelling framework that has been used in the development of JSDMs is explicitly linear, which it straightforward to write down models with extra effects. Although fitting these models may not be easy, software has been developed to fit these models (Ovaskainen, Tikhonov, Norberg, et al. 2017; Niku et al. 2019), and extensions beyond these can be developed using flexible Bayesian packages such as JAGS (Plummer 2021) and Nimble (de Valpine et al. 2017).

The JSDM approach also helps with incorporating better sampling models (e.g. Beissinger et al. 2016; Björk, Hui, et al. 2018; Tobler et al. 2019), so that issues like pseudo-replication can be handled by properly incorporating their effects into the model. For example, multiple traps or visits at a site can be treated as replicate samples from the same community: with binary data this can be used in an occupancy model (Tobler et al. 2019), but when the data estimate abundance (e.g. through counts), the repeated samples can be used to estimate the amount of sampling error. These models can be extended to include observation-level covariates, in the same way that they are in occupancy models.

Another area where ordination can follow methods developed in the JSDM world is the use of temporally explicit models. Because the approach is based on an explicit model, it can include a temporal autocorrelation (e.g. Björk, O’Hara, et al. 2018; Ovaskainen, Tikhonov, Dunson, et al. 2017). This is preferable to the approach that classical ordination takes, where the order of the years is ignored in the ordination, although it can be incorporated into the graphical presentation (e.g. Blanchette and Pearson 2013). This links the temporal changes more directly to models of community dynamics, as it is a multi-species Gompertz model where environmental effects on growth rates can be incorporated (e.g. Mutshinda, O’Hara, and Woiwod 2011).

## Hierarchical Ordination; a unifying framework for drivers of community processes

The GLLVM framework is flexible enough to be developed in several directions. Site scores have already been modelled as a function of covariates, through a concurrent GLLVM (van der Veen, Hui, Hovstad, and O’Hara 2021), and also as a spatial field (allowing for spatial autocorrelation), in a spatial factor analysis (e.g. Thorson et al. 2015). But species effects, such as traits or phylogenetic effects, could be modelled instead, simply by transposing the data matrix. Ordination can thus be linked to trait-based analyses.

In the fixed effects, estimating the interactions between site and species covariates are the “fourth corner problem”. This can be parameter heavy when there are several covariates (e.g. see Niku et al. 2021 in the context of GLLVMs). One possible approach is to extend the constrained ordination idea, so that both site scores and species loadings are modelled further. This leads us to the idea of what we have called hierarchical ordination, an extension of double constrained ordination (Cajo J. F. ter Braak, Šmilauer, and Dray 2018). As before we have

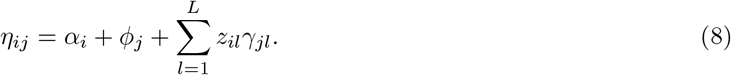

But now we extend the models for both *z*_*il*_ and *γ*_*jl*_:

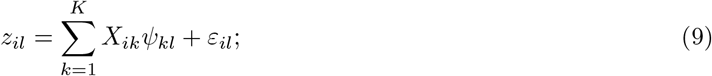

And

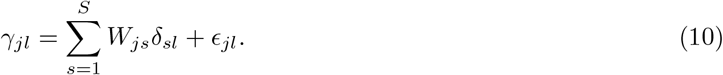

Because both *z*_*il*_ and *γ*_*jl*_ can be written as sums of other terms, it is straightforward model them both hierarchically, as we would for any other hierarchical model. For example, a spatial effect can be added to the site scores, similar to the spatial factor analysis idea, but trait and phylogenetic effects can also be added to the species loadings. The method developed by Cajo J. F. ter Braak, Šmilauer, and Dray (2018) is similar, but they assume 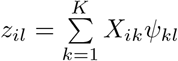 and 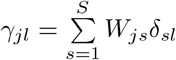, i.e. the site scores and species effects are fully determined by the covariates.

We can also standardise *z*_*il*_ and *γ*_*jl*_ so each has a variance of one, and then introduce a matrix of scaling parameters **Λ**. The main diagonal of this matrix can be interpreted in the same way as an singular values in classical ordination: it summarises how much of the variation can be explained by that latent variable. Typically all of the values off the main diagonal would be zero, although this is not necessary: off-diagonal terms would mean that the latent variables are correlated. Consequently, hierarchical ordination is more similar in form to classical ordination methods than existing model-based ordination methods.

Expanding the model, we get

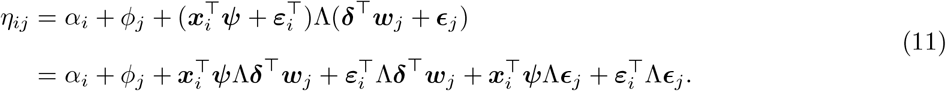

This model acts as an reduced version for the fourth corner latent variable model because it incorporates trait by environment interactions through the latent variables. Going through the ordination terms in, scalar notation, we have

- **Λ**: the variance of the latent variable (on the link scale), similar to singular values i classical ordination
- 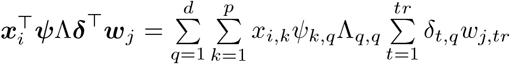: the fourth-corner term approximated in reduced rank, or a double constrained ordination
- 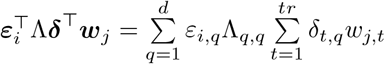: a constrained ordination where the species loadings are constrained by traits alone
- 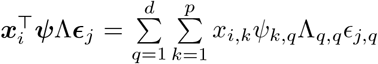: a constrained ordination with site effects determined by environmental covariates
- 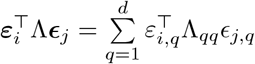: a residual ordination

Each of which would have a different place in the analysis of co-occurrence patterns in ecological communities. Of course, these terms can also include structured random effects, e.g. spatial or phylogenetic.

Another way to look at this model is that it can model the change in associations between species with the environment (Tikhonov et al. 2017; Perrin et al. 2021). This comes from the fourth corner term, which models the interaction between traits and environment, so plays the same role as equation (7).

One advantage of having an explicit hierarchical ordination framework is that visualisation and interpretation can be based on current ordination methods, for example ordination diagrams can be drawn for both site and species effects. Thus the advantages of ordination as a way of summarising correlation, and the effects of covariates on the correlations, are retained with this model.

There may be problems where a single hierarchical ordination is not sufficient. For example, an additional species-specific covariate effect may be needed, leading to this model:

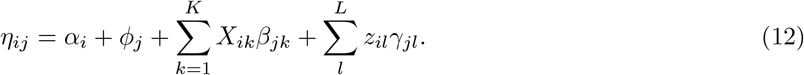

Potentially, more than one ordination could also be used (Björk, Hui, et al. 2018), so the framework is flexible, at the cost (both computationally and in terms of interpretation) of adding extra latent levels. Using a single ordination, with effects on either species or sites, we simplify the model into manageable parts. The choice of whether to use one or two (or more!) sets of ordination will depend on whether a single ordination is reasonable ecologically, and whether multiple ordinations are feasible computationally. Whilst the complexity of additional ordinations is attractive, the price is that more data will be needed to estimate the parameters of the model, and the fitting will be more difficult.

#### An example of hierarchical ordination

We demonstrate hierarchical ordination with data originally collected by Ribera et al. (2001). It consists of abundance observations of 68 ground beetle species on 87 sites in Scotland, along with 17 environmental variables and 20 traits. The aim to is determine how the environmental variables and traits affect composition in the ecological community, including whether they interact. With so many traits and environmental variables, hierarchical ordination is an attractive approach to controlling the variation.

The model is described in the Supporting Information. Briefly, we model the counts as following a negative-binomial distribution with a log link, two latent variables and linear models for the site scores (with the 17 environmental variables) and species effects (20 traits).

Biplots of the species and site effects are in Fig. 1. The traits and the environment explain 47 and 75 percent of the variation explained by the model. From the 10 estimated latent variables, the first two explain 60 percent of the variation, of which the environmental variables explain 1 percent and 11 percent of the variation in the site scores of the first and second latent variables respectively, and the traits explain 12 percent and 13 percent of the variation in the species effects, indicating that the dominant ecological gradients cannot be represented well by the covariates. Most notably, the latent variables with the largest variance explained by the predictors are the third and fifth for environment and traits, respectively. We can see that the first axis is largely a management gradient, and (in essence) the maximum width and having black legs (CLG2) are the main traits that explain abundance variation. We note that both of these may be phylogenetically conserved.

**Figure 1:**
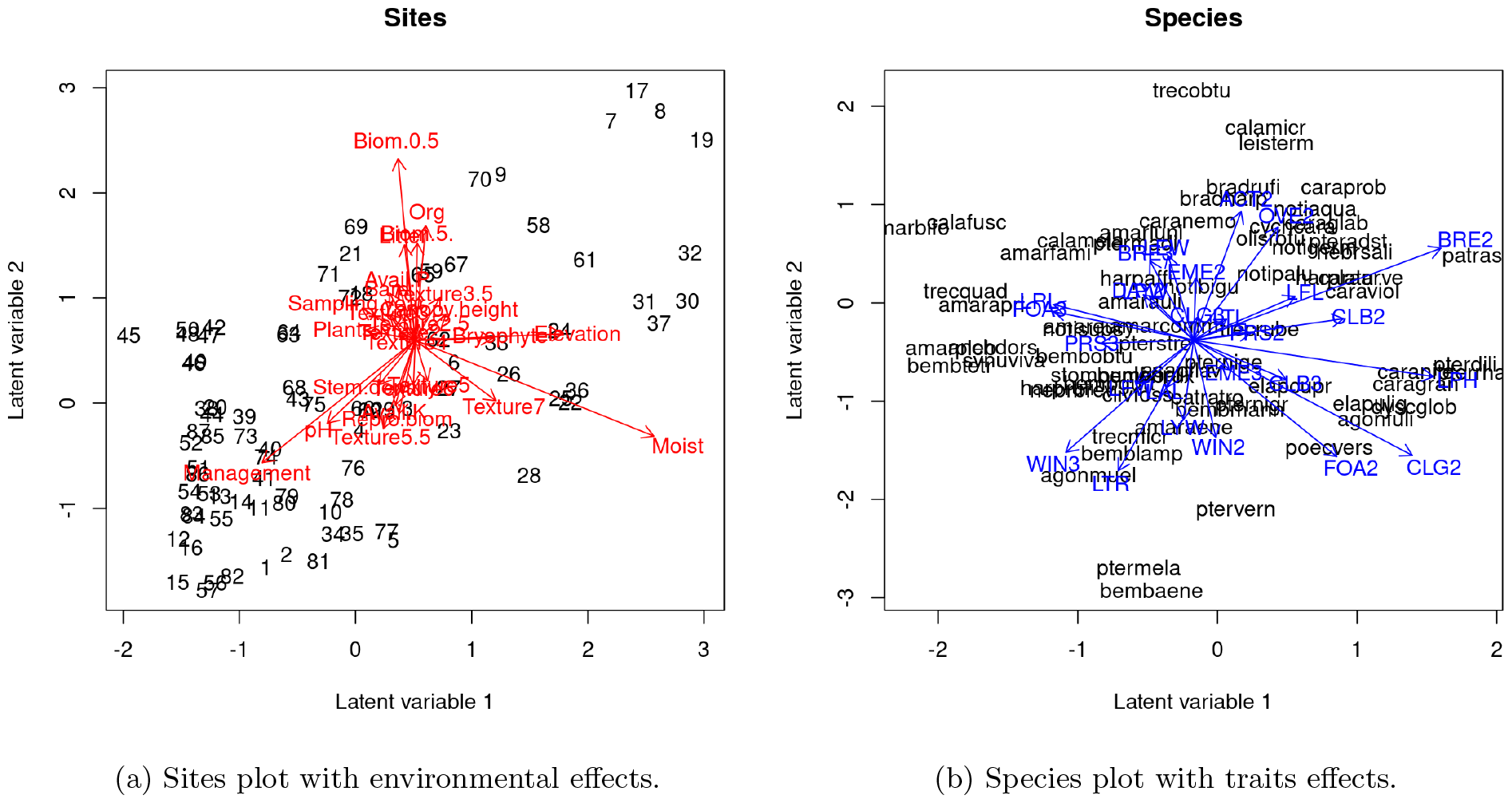
Visualization of the hierarchical ordination for the beetle data. Note, that the environmental effects could have been plotted in the species plot, or the trait effects in the sites plot, instead.

## Discussion

We have discussed how ordination and JSDMs have converged onto a similar set of models, GLLVMs, providing a common framework that is available to both groups of ecologists. The differences between the two approaches largely came from scale and the questions being asked, but the models are flexible enough to handle these. Having a single set of models should help to unify these different approaches to community ecology, and the link to temporal models should improve connections to ecological theory, efficiently integrating data into the frameworks that are being developed.

A model-based approach allows for a lot of flexibility, as we can see in our overview. One downside of this flexibility is the possibility that the model becomes too complex to be interpretable. With a large number of species and sites, it is easy to develop models that ask for all of this information to be used. Our suggestion of a hierarchical approach to ordination tries to reduce this complexity by making the parameters either site- or species- specific. Interactions between the two sets of parameters are made through the latent variables. This is one simplification of the model, so the response of a community to changes in the environment are measured by how the site scores change, and then how this affects species, through their loadings. This shifts focus from individual species or sites to the community, and the gradients.

It is one thing to write down a model, but another to fit it. For “power users” flexible software, like Nimble (de Valpine et al. 2017), which takes advantage of the flexibility and simplicity of the BUGS language, can be used to develop and extend these models. The example shown here was fitted with Nimble, and the code is available in the Supporting Information. But for most ecologists it would be better to have bespoke software, which would mean developing packages like HMSC (Ovaskainen, Tikhonov, Norberg, et al. 2017) and gllvm (Niku et al. 2019) to fit models in the full framework.

We would not want to suggest that every ordination should use this framework: there will be times when the question being asked requires a different model. However, we feel that our framework is sufficiently flexible for many problems, and will build on both the modelling strengths developed from JSDMs and the visualisation and interpretation provided by the ordination world^1^. If extensions are needed, we would suggest that our model provides a starting point, both in terms of writing down the extended model and also in justifying why such an extension is needed. For example, when Björk, Hui, et al. (2018) used two ordinations to model two levels of sampling, a natural question is whether it would have been better to use one ordination, with sample and host species levels in the site effects, and with the same species loading. The answer to this question is not just one of modelling, it is also ecologically informative, i.e. about the extent to which species were responding to hosts, or to site effects.

Though the modeling framework suggested here is statistical in nature, this approach can serve to provide new insight into ecological theories on community assemblages. Further extensions of the framework towards spatiotemporal processes can serve to develop a theoretical model for how species interact in space and time, and how species-specific properties relate to site-specific properties, as in the case of functional traits.

The modelling of multispecies communities is being changed by the application of more sophisticated statistical techniques. Here we are suggesting that they can unify different fields, as they are modelling similar processes, and so can open up the fields to new questions. This is done both by moving ideas from one field to another (e.g. using ordination plots in JSDMs), and also by opening up new ways of analysing the data, with flexible models that can handle the problems associated with having many sites and species. To quote one of the world’s great philosophers, let’s go exploring (Watterso 1995).

## Supporting information

Hierarchical Ordination Vignette

## Author Contributions

Usually a manuscript with a student and their supervisor comes about when the supervisor has a brilliant idea, but doesn’t have time, so the student has to do all of the hard work, trying to find out if the idea actually works and doing the writing. In this manuscript, the roles were reversed. Neither author is totally sure how it happened.

## Footnotes

1 a world that is, of course, only two dimensional

## References

Batterink, Marten. 1983. “Een Vergelijkend Vegetatiekundig Onderzoek Naar de Typologie En Invloeden van Het Beheer van 1973 Tot 1982 in de Duinweilanden Op Terschelling.” PhD thesis, Landbouwhogeschool.

Beissinger, Steven R., Kelly J. Iknayan, Gurutzeta Guillera-Arroita, Elise F. Zipkin, Robert M. Dorazio, J. Andrew Royle, and Marc Kéry. 2016. “Incorporating Imperfect Detection into Joint Models of Communities: A Response to Warton Et Al.” Trends in Ecology & Evolution 31 (10): 736–37. 10.1016/j.tree.2016.07.009.

Björk, Johannes R., Francis K. C. Hui, Robert B. O’Hara, and Jose M. Montoya. 2018. “Uncovering the Drivers of Host-Associated Microbiota with Joint Species Distribution Modelling.” Molecular Ecology 27 (12): 2714–24. 10.1111/mec.14718.

Björk, Johannes R., Robert B. O’Hara, Marta Ribes, Rafel Coma, and M. José Montoya. 2018. “The Dynamic Core Microbiome: Structure, Dynamics and Stability.” bioRxiv : The Preprint Server for Biology. 10.1101/137885.

Blanchette, M. L., and R. G. Pearson. 2013. “Dynamics of Habitats and Macroinvertebrate Assemblages in Rivers of the Australian Dry Tropics.” Freshwater Biology 58 (4): 742–57. 10.1111/fwb.12_080.

Braak, Cajo J F ter, Petr Šmilauer, and Stéphane Dray. 2018. “Algorithms and Biplots for Double Constrained Correspondence Analysis.” Environmental and Ecological Statistics 25 (2): 171–97.

Braak, Cajo J. F. ter. 1985. “Correspondence Analysis of Incidence and Abundance Data: Properties in Terms of a Unimodal Response Model.” Biometrics. Journal of the International Biometric Society 41 (4): 859–73. http://www.jstor.org/stable/2530959.

Braak, Cajo J. F. ter. 1986. “Canonical Correspondence Analysis: A New Eigenvector Technique for Multivariate Direct Gradient Analysis.” Ecology 67 (5): 1167–79. 10.2307/1938672.

Bray, J Roger, and John T Curtis. 1957. “An Ordination of the Upland Forest Communities of Southern Wisconsin.” Ecological Monographs 27 (4): 326–49.

Clark, James S, Alan E Gelfand, Christopher W Woodall, and Kai Zhu. 2014. “More Than the Sum of the Parts: Forest Climate Response from Joint Species Distribution Models.” Ecological Applications 24 (5): 990–99.

Gabriel, K. R. 1971. “The Biplot Graphic Display of Matrices with Application to Principal Component Analysis.” Biometrika 58 (3): 453–67. 10.1093/biomet/58.3.453.

Gower, J. C. 1966. “Some Distance Properties of Latent Root and Vector Methods Used in Multivariate Analysis.” Biometrika 53 (3-4): 325–38. 10.1093/biomet/53.3-4.325.

Hui, Francis K. C., Sara Taskinen, Shirley Pledger, Scott D. Foster, and David I. Warton. 2015. “Model-Based Approaches to Unconstrained Ordination.” Methods in Ecology and Evolution 6 (4): 399–411. 10.1111/2041-210X.12236.

Jamil, Tahira, and Cajo J F Ter Braak. 2013. “Generalized Linear Mixed Models Can Detect Unimodal Species-Environment Relationships.” PeerJ 1 (July): e95.

Kissling, W. D., Carsten F. Dormann, Jürgen Groeneveld, Thomas Hickler, Ingolf Kühn, Greg J. McInerny, José M. Montoya, et al. 2012. “Towards Novel Approaches to Modelling Biotic Interactions in Multispecies Assemblages at Large Spatial Extents.” Journal of Biogeography 39 (12): 2163–78. 10.1111/j.1365-2699.2011.02663.x.

Mutshinda, Crispin M., Robert B. O’Hara, and Ian P. Woiwod. 2011. “A Multispecies Perspective on Ecological Impacts of Climatic Forcing.” Journal of Animal Ecology 80 (1): 101–7. 10.1111/j.1365-2656.2010.01743.x.

Niku, Jenni, Francis K. C. Hui, Sara Taskinen, and David I. Warton. 2019. “Gllvm - Fast Analysis of Multivariate Abundance Data with Generalized Linear Latent Variable Models in R.” Methods in Ecology and Evolution 10: 2173–82.

Niku, Jenni, Francis K. C. Hui, Sara Taskinen, and David I. Warton. 2021. “Analyzing Environmental-Trait Interactions in Ecological Communities with Fourth-Corner Latent Variable Models.” Environmetrics (London, Ont.) 32 (6): e2683. 10.1002/env.2683.

O’Hara, R. B., and M. J. Sillanpää. 2009. “A Review of Bayesian Variable Selection Methods: What, How and Which.” Bayesian Analysis 4 (1): 85–117. 10.1214/09-BA403.

Ovaskainen, Otso, Gleb Tikhonov, David Dunson, Vidar Grøtan, Steinar Engen, Bernt-Erik Sæther, and Nerea Abrego. 2017. “How Are Species Interactions Structured in Species-Rich Communities? A New Method for Analysing Time-Series Data.” Proceedings of the Royal Society B: Biological Sciences 284 (1855): 20170768. 10.1098/rspb.2017.0768.

Ovaskainen, Otso, Gleb Tikhonov, Anna Norberg, Guillaume F. Blanchet, Leo Duan, David Dunson, Tomas Roslin, and Nerea Abrego. 2017. “How to Make More Out of Community Data? A Conceptual Framework and Its Implementation as Models and Software.” Ecology Letters 20 (5): 561–76. 10.1111/ele.12757.

Perrin, Sam Wenaas, Bert van der Veen, Nick Golding, and Anders Gravbrøt Finstad. 2021. “Modelling Temperature-Driven Changes in Species Associations Across Freshwater Communities.” Global Change Biology n/a (n/a). 10.1111/gcb.15888.

Plummer, Martyn. 2021. Rjags: Bayesian Graphical Models Using MCMC. Manual. https://CRAN.R-project.org/package=rjags.

Pollock, Laura J., Reid Tingley, William K. Morris, Nick Golding, Robert B. O’Hara, Kirsten M. Parris, Peter A. Vesk, and Michael A. McCarthy. 2014. “Understanding Co-Occurrence by Modelling Species Simultaneously with a Joint Species Distribution Model (JSDM).” Methods in Ecology and Evolution 5 (5): 397–406. 10.1111/2041-210X.12180.

Skrondal, Anders, and Sophia Rabe-Hesketh. 2004. Generalized Latent Variable Modeling: Multilevel, Longitudinal, and Structural Equation Models. Chapman and Hall/CRC.

Thorson, James T., Mark D. Scheuerell, Andrew O. Shelton, Kevin E. See, Hans J. Skaug, and Kasper Kristensen. 2015. “Spatial Factor Analysis: A New Tool for Estimating Joint Species Distributions and Correlations in Species Range.” Methods in Ecology and Evolution 6 (6): 627–37. 10.1111/2041-210X.12359.

Tikhonov, Gleb, Nerea Abrego, David Dunson, and Otso Ovaskainen. 2017. “Using Joint Species Distribution Models for Evaluating How Species-to-Species Associations Depend on the Environmental Context.” Methods in Ecology and Evolution 8 (4): 443–52. 10.1111/2041-210X.12723.

Tikhonov, Gleb, Li Duan, Nerea Abrego, Graeme Newell, Matt White, David Dunson, and Otso Ovaskainen. 2020. “Computationally Efficient Joint Species Distribution Modeling of Big Spatial Data.” Ecology 101 (2): e02929.

Tobler, Mathias W., Marc Kéry, Francis K. C. Hui, Gurutzeta Guillera-Arroita, Peter Knaus, and Thomas Sattler. 2019. “Joint Species Distribution Models with Species Correlations and Imperfect Detection.” Ecology 100 (8): e02754. 10.1002/ecy.2754.

Tredennick, Andrew T., Giles Hooker, Stephen P. Ellner, and Peter B. Adler. 2021. “A Practical Guide to Selecting Models for Exploration, Inference, and Prediction in Ecology.” Ecology 102 (6): e03336. 10.1002/ecy.3336.

Valpine, Perry de, Daniel Turek, Christopher Paciorek, Cliff Anderson-Bergman, Duncan Temple Lang, and Ras Bodik. 2017. “Programming with Models: Writing Statistical Algorithms for General Model Structures with NIMBLE.” Journal of Computational and Graphical Statistics 26 (2): 403–13. 10.1080/10618600.2016.1172487.

Veen, Bert van der, Francis K. C. Hui, Knut A. Hovstad, and Robert B. O’Hara. 2021. “Model-Based Ordination with Constrained Latent Variables.” bioRxiv : The Preprint Server for Biology. 10.1101/2021.10.11.463884.

Veen, Bert van der, Francis K. C. Hui, Knut A. Hovstad, Erik B. Solbu, and Robert B. O’Hara. 2021. “Model-Based Ordination for Species with Unequal Niche Widths.” Methods in Ecology and Evolution 12 (7): 1288–1300. 10.1111/2041-210X.13595.

Warton, David I., F. Guillaume Blanchet, Robert B. O’Hara, Otso Ovaskainen, Sara Taskinen, Steven C. Walker, and Francis K. C. Hui. 2015. “So Many Variables: Joint Modeling in Community Ecology.” Trends in Ecology & Evolution 30 (12): 766–79. 10.1016/j.tree.2015.09.007.

Watterso, Bill. 1995. “Final Calvin and Hobbes - Last Comic - by Bill Watterson for December 31, 1995 Calvin and Hobbes Comic Strip for December 31, 1995.” 1995. https://www.gocomics.com/calvinandho bbes/1995/12/31.

Wilkinson, David P., Nick Golding, Gurutzeta Guillera-Arroita, Reid Tingley, and Michael A. McCarthy. 2021. “Defining and Evaluating Predictions of Joint Species Distribution Models.” Methods in Ecology and Evolution 12 (3): 394–404. 10.1111/2041-210X.13518.

Wisz, Mary Susanne, Julien Pottier, W. Daniel Kissling, Loïc Pellissier, Jonathan Lenoir, Christian F. Damgaard, Carsten F. Dormann, et al. 2013. “The Role of Biotic Interactions in Shaping Distributions and Realised Assemblages of Species: Implications for Species Distribution Modelling.” Biological Reviews 88 (1): 15–30. 10.1111/j.1469-185X.2012.00235.x.

