## Supplementary material for "Hierarchical Ordination, A unifying framework for drivers of community processes": Hierarchical Ordination Vignette


### Hierarchical ordination

#### 2024-01-08

### An Example of Hierarchical Ordination

This document describes fitting a hierarchical ordination to data,
including code and the details that are needed. This is all available on
GitHub,
except for the MCMC output, which is (at the moment) on
Dropbox.

#### The data

The data is available on figshare,
but was originally collected by Ribera
*et al.* (2001), and reanalyzed by Niku
*et al.* (2021). It includes abundance observations of 68
ground beetles 87 sites in Scotland, along with 17 environmental
variables and 20 traits. The aim to is determine how the environmental
variables and traits affect composition in the ecological community,
including whether they interact. The methodological question is how does
hierarchical ordination help.

The abundances, traits and environment are each stored in a different
matrix. First we load the data and set some constants:

```
library(nimble)
library(nimbleHMC)
library(coda)
library(lattice)

# Response data
Y <- t(read.csv("Y.csv"))
colnames(Y) <- Y[2,]
Y<-Y[-c(1:2),-c(1,70:71)]
Y <- as.data.frame(apply(Y,2,as.integer))

# Environmental predictors
X <- read.csv("X.csv")[,-c(1:5)]
X <- as.data.frame(apply(X,2,as.numeric))
X$Sampling.year <- X$Sampling.year - min(X$Sampling.year)
X$Texture <- as.factor(X$Texture)

# Traits
TR  <- read.csv("TR.csv")
row.names(TR) <- TR$SPECIES
TR <- TR[,-c(1:3)]
# Traits to categorical
# Removing question marks, not ideal
TR[,c("CLG","CLB","WIN","PRS","OVE","FOA","DAY","BRE","EME","ACT")] <- apply(TR[,c("CLG","CLB","WIN","PRS","OVE","FOA","DAY","BRE","EME","ACT")],2,function(x)as.factor(gsub("\\?.*","",x)))

# Data standardization
X <- scale(model.matrix(~.,X))[,-1] # environmental variables
TR <- scale(model.matrix(~.,TR))[,-1] # species traits

# Constants
NSites <- nrow(Y) # number of sites
NSpecies <- ncol(Y) # number of species
NTraits <- ncol(TR) # number of traits
NEnv <- ncol(X) # number of environmental predictors

# create data lists for Nimble
dat <- list(Y = Y, X = as.matrix(X), TR = as.matrix(TR))
consts <- list(NSites = NSites, NEnv = NEnv, NTraits = NTraits, NSpecies = NSpecies)
```

#### The Model

The data are counts of each species so we assume they follow a
Poisson distribution with a log link function, as we would do in a
standard generalised linear model. We assume that each species has a
different mean abundance (i.e. for each species \(j\) we have a different intercept \(\beta\_{0j}\)), and model the rest of the
variation with a hierarchical ordination. This gives the following mean
model on the link scale (with linear predictor \(\eta\_{ij}\)):

\[
\eta\_{ij} = \beta\_{0j} + \boldsymbol{z}\_i^\top \boldsymbol{\Sigma}
\boldsymbol{\gamma}\_j.
\]

As with any ordination, \(\boldsymbol{z}\) and \(\boldsymbol{\gamma}\_j\) are the site scores
and species loadings, and the columns of \(\boldsymbol{Z}\) are the latent variables
(which holds the site scores as the rows). Here we assume that they each
have a variance of one, so that \(\boldsymbol{\Sigma}\) holds the variation
of the latent variables: it will typically be a diagonal matrix, and if
any of the terms on the diagonal are close to zero, this suggests that
latent variable has (almost) no effect. It thus provides a
straightforward summary of the relative importance of that latent
variable, and is similar to the square root of eigenvalues (singular
values) in an eigenanalysis.

If we stopped here, this would be a standard Generalized Linear
Latent Variable Model. But, here we want to model both \(\boldsymbol{z}\_i\) and \(\boldsymbol{\gamma}\_j\), i.e. go from
modelling model groups of species responding in similar ways to sites to
modelling how the traits of the species affect their responses to the
environment.

As a simplification, we can think about this as simply a regression
of \(\boldsymbol{z}\_i\) (the site
effects) against the environmental variables and \(\boldsymbol{\gamma}\_j\) against the traits.
In reality, it is more complicated, because \(\boldsymbol{z}\_i\) and \(\boldsymbol{\gamma}\_j\) are estimated in
the model, so their uncertainty needs to be propagated.

Some more mathematical details are hidden away here, for those who are
interested.

We denote the abundance at site \(i = 1
\dots n\) for species \(j = 1 \dots
p\) as \(Y\_{ij}\), or as a
matrix as \(\boldsymbol{Y}\). The
environmental variables are \(x\_{ik}\)
for the \(k = 1 \ldots K\) predictors
(or \(\boldsymbol{X}\) as a matrix),
and \(t = 1\ldots T\) traits as \(\boldsymbol{TR}\). We then assume

\[Y\_{ij} \sim
\text{Pois}(\lambda\_{ij}),\]

with

\[\text{log}(\lambda\_{ij}) =
\eta\_{ij}.\]

Consequently, \(\eta\_{ij}\) is the
linear predictor, which we further model with the hierarchical
ordination:

\[
\eta\_{ij} = \beta\_{0j} + \boldsymbol{z}\_i^\top \boldsymbol{\Sigma}
\boldsymbol{\gamma}\_j.
\]

We then model \(\boldsymbol{z}\_i\)
(as in van
der Veen *et al.* (2023)) and \(\boldsymbol{\gamma}\_j\) hierarchically:

\[
\boldsymbol{z}\_i = \boldsymbol{B}^\top\boldsymbol{x}\_i +
\boldsymbol{\epsilon}\_i
\] and

\[
\boldsymbol{\gamma}\_j = \boldsymbol{\omega}^\top\boldsymbol{TR}\_{j} +
\boldsymbol{\varepsilon}\_j
\] where:

- \(x\_{ik}\) is the \(k^{th}\) predictor (i.e. environmental
  effect) at site \(i\)
- \(\boldsymbol{B}\) with entry \(b\_{kq} \sim \mathcal{N}(0,1)\) is the
  effect of the \(k^{th}\) predictor on
  the site score for the \(q^{th} = 1\ldots
  d\) latent variable
- \(\boldsymbol{\epsilon}\_i\) with
  entry \(\epsilon\_{iq} \sim \mathcal{N}(0,
  \sigma^2\_q)\) is a vector of residuals for the unexplained part
  of the site score
- \(TR\_{jt}\) is the \(t^{th}\) predictor (i.e. trait) for species
  \(j\)
- \(\boldsymbol{\omega}\_t\) with
  entry \(\omega\_{tq}\) is the effect of
  the \(t^{th}\) trait on the species
  loading for the \(q^{th}\) latent
  variable
- \(\boldsymbol{\varepsilon}\_j\) with
  entry \(\varepsilon\_{jq} \sim \mathcal{N}(0,
  \delta^2\_q)\) is a vector of residuals for the unexplained part
  of the species loading

Note that the predictors are all standardized to zero mean and unit
variance. We additionally place exponential priors with rate parameter
one on all the scale parameters.

### Implementation

We fit the model with the *Nimble* *R*-package. We
start with a single dimension for simplicity, so that we can show the
steps needed.

Functions needed for this vignette are here.


Functions for MCMC are here.

```
# Function to run one chain: it can be done with HMC or other MCMC algorithms.
# "block" can be used to specify blocking structures for the slice sampler
# "slice" can be used to specify parameters on which to apply univariate Metropolis-Hastings
# parameters that are not in "block" or "slice" are HMCed
# Function to run a single chain
RunOneChain <- function(seed, dat, code, inits, consts, ToMonitor=NULL, 
                        Nburn=5e3, NIter=5.5e4, Nthin=10, block = NULL, slice = NULL, ...) {
  require(nimble)
  require(nimbleHMC)
  AllSamplers <- HMCsamplers <- c('epsilonSTAR', 'varepsilonSTAR', 'beta0', 'OSTAR', 'BSTAR',
                  'sd.SiteSTAR', 'sd.SpeciesSTAR','xi', 'phi')
  
  if(!is.null(block)){
  HMCsamplers <- HMCsamplers[!HMCsamplers%in%unique(gsub("\\s*\\[[^\\)]+\\]","",c(unlist(block),unlist(slice))))]
  }
  if(is.null(ToMonitor)) {
    ToMonitor <- c("beta0", "sd.SpeciesSTAR", "sd.SiteSTAR", "sd.LV", "BSTAR", "OSTAR", "B","O",
                   "epsilonSTAR", "varepsilonSTAR", "xi", "phi")
  }
  mod <- nimbleModel(code = code, name = "HO", inits = inits(consts), constants = consts, data = dat, buildDerivs = TRUE)
  model <- compileNimble(mod)
  
  # Do HMC
    conf <- configureHMC(model, nodes = HMCsamplers, monitors = ToMonitor, print = FALSE, 
                         control=list(nwarmup=Nburn))
    if(!is.null(block)) { 
    if(is.list(block)){
    # Use a slice everything that not being HMCed
      lapply(block, conf$addSampler, type = "AF_slice", sliceAdaptFactorInterval = Nburn)
    }else{
      # Use a slice everything that not being HMCed
      sapply(block, conf$addSampler, type = "AF_slice", sliceAdaptFactorInterval = Nburn)
    }
    }
    
    if(!is.null(slice)) { 
    if(is.list(slice)){
    # Use a slice everything that not being HMCed
      lapply(slice, conf$addSampler, type = "slice")
    }else{
      # Use a slice everything that not being HMCed
      sapply(slice, conf$addSampler, type = "slice")
    }
    }
    
  mcmc <- buildMCMC(conf)
  cmcmc <- compileNimble(mcmc, project = model)
  res <- runMCMC(cmcmc,  niter=NIter, nburnin = Nburn, thin=Nthin, 
                 nchains = 1, samplesAsCodaMCMC = TRUE)
  return(res)
}

# Function to run chains in  parallel
ParaNimble <- function(NChains, ...) {
  opts <- list(...)
  if(!is.null(opts$seeds) && (length(opts$seeds) == NChains)){
    seeds <- opts$seeds
    opts <- opts[-which(names(opts)=="seeds")]
  }else{
    seeds <- 1:NChains
  }
    
  require(parallel)
  nimble_cluster <- makeCluster(NChains)
  clusterExport(nimble_cluster, "nimQR.u")
  samples <- parLapply(cl = nimble_cluster, X = seeds, ...)
  stopCluster(nimble_cluster)

  # Name the chains in the list
  chains <- setNames(samples,paste0("chain", 1:length(samples)))
  chains
}

# Function to create list of names for parameters to use a block slice sampler for

MakeBlockList <- function(consts, LVwise = TRUE){
# builds list for slice AF sampling
# with LVwise = TRUE we are LV-wise blocking B and epsilon, and O and varepsilon
# otherwise, LVs are jointly blocked
if(LVwise & consts$nLVs>1){
blockList <- c(sapply(1:consts$nLVs,
       function(x,consts){
         c(paste0("BSTAR[", 1:consts$NEnv,", ",x, "]"),
           paste0("epsilonSTAR[", 1:consts$NSites,", ",x, "]"))
         }
       ,
         consts = consts,simplify=F),
       sapply(1:consts$nLVs,
       function(x,consts){
         c(paste0("OSTAR[", 1:consts$NTraits,", ",x, "]"),
           paste0("varepsilonSTAR[", 1:consts$NSpecies,", ",x, "]"))
         }
       ,
         consts = consts,simplify=F)
       )
}else{
  blockList = list(c("BSTAR","epsilonSTAR"),c("OSTAR","varepsilonSTAR"))
}
blockList
}
```


Functions for processing the MCMC are here.

```
GetMeans <- function(summ, name, d) {
  var <- summ$statistics[grep(paste0('^', name), rownames(summ$statistics)),"Mean"]
  matrix(var,ncol=d)
}

# Utility Function to get logical indicators for if names contains a string in v
GetInds <- function(v, names) {
  if(length(v)==1) {
    res <- grep(v, names)
  } else {
    res <- c(unlist(sapply(v, grep, x=names)))
  }
  res
}

# Function to swap signs of all variables varinds to have same sign as vartosign.
ReSignChain <- function(chain, varinds, vartosign) {
  #    MeanSign <- sign(mean(chain[  ,s])) # Might need this
  res <- t(apply(chain, 1, function(x, vs, vi) {
    Names <- names(x)[vi]
    if(any(grepl(",", Names))) {
      lvind <- gsub(".*, ", "", Names)
    } else {
      lvind <- seq_along(vs)
    }
    x[vi] <- x[vi]*sign(x[vs[lvind]])
    x
  }, vs=vartosign, vi=varinds))
  as.mcmc(res)
}

# Function to post-process chains to swap signs.
postProcess <- function(Chains, VarsToProcess, VarsToSwapBy = NULL, VarToSign=NULL, print=FALSE, rule = 2) {
  if(is.null(VarToSign)) VarToSign <- VarsToProcess
  SignInd <- GetInds(VarToSign, colnames(Chains[[1]]))
  # Get indicators for all variables to have their signs changed
  ProcessInds <- GetInds(VarsToProcess, names = colnames(Chains[[1]]))
  
  # Check if > 1 LV
  SeveralLVs <- any(grepl(",", colnames(Chains[[1]])[SignInd]))
  LV <- gsub(".*, ", "", colnames(Chains[[1]])[SignInd])
  
  if(rule==1){
  # Calculate variance of mean of indicator of sign: 
  # hopefully largest is variable with most sign swapping (i.e. )
  Signs <- as.data.frame(lapply(Chains, function(mat, ind) {
    colMeans(mat[,ind]>0)
  }, ind=SignInd))
  
  VarSign <- apply(Signs, 1, var)
  
  if(SeveralLVs) {
    LV <- gsub(".*, ", "", colnames(Chains[[1]])[SignInd])
    #Chose variables who's sign will be used to swap other signs
    if(is.null(VarsToSwapBy)){
      VarsToSwapBy <- sapply(unique(LV), function(lv, vs) {
      vv <- vs[grep(lv, names(vs))]
      nm <- names(which(vv==max(vv)))
      if(length(nm)>1) nm <- nm[1] # probably something more sophisticated is better
      nm
    }, vs = VarSign, simplify = TRUE)
    }
    
  } else { # only 1 LV
    if(is.null(VarsToSwapBy)){
    #Chose variables who's sign will be used to swap other signs
    VarsToSwapBy <- names(which(VarSign==max(VarSign)))[1]
    }
  }
  
  }else if(rule==2){
    require(mousetrap)
    
    jointChains <- do.call(rbind, Chains)
    # bimodality score
    bms<-apply(jointChains[,SignInd],2,bimodality_coefficient)
    # find maximum bimodality score
    if(SeveralLVs){
    lstSgn <- sapply(unique(LV),function(lv)which.max(bms[grep(lv,names(bms))]),simplify=F)
    # formatting
    names(lstSgn) <- NULL
    VarsToSwapBy <- names(unlist(lstSgn))
    names(VarsToSwapBy) <- paste0(sort(unique(LV)))
    }else{
    lstSgn <- which.max(bms)
    # formatting
    VarsToSwapBy <- names(lstSgn)
    names(VarsToSwapBy) <- "1]"
    }
  }
  
  if(print) message(paste0("Swapping by ", paste(VarsToSwapBy, collapse=", ")))
  chains.sgn <- lapply(Chains, ReSignChain, vartosign=VarsToSwapBy, varinds=ProcessInds)

  as.mcmc.list(chains.sgn)
}

# Function to convert a variable in an MCMC chain that should be a matrix from 
# a vector to the right matrix
ChainToMatrix <- function(ch, name) {
  v <- ch[grep(paste0("^", name, "\\["), names(ch))]
  l.row <- as.numeric(gsub(".*\\[", "", gsub(",.*", "", names(v))))
  l.col <- as.numeric(gsub(".*, ", "", gsub("\\]", "", names(v))))
  mat <- matrix(0, nrow=max(l.row), ncol=max(l.col))
  mat[l.row+max(l.row)*(l.col-1)]<-v
  mat
}

# Function to convert a variable in an MCMC chain that should be a vector to a diagonal matrix
ChainToDiag <- function(ch, name) {
  v <- ch[grep(paste0("^", name, "\\["), names(ch))]
  mat <- diag(v)
  mat
}

RescaleVars <- function(vec, ScaleBy, SDTorescale,ToRescale) {
  if(!any(c("z","gamma")%in%ScaleBy))stop("Not a valid choice.")
  if("sd.LV"%in%SDTorescale)stop("Not a valid choice.")
  vec2 <- vec
  # get scale from z or gamma
  ScBy <- ChainToMatrix(vec, ScaleBy)
  SD <- apply(ScBy, 2, sd)
  
  for(nm in unique(c(ScaleBy, ToRescale))) {
      Sc.x <- ChainToMatrix(vec, nm)
      Sc.x.sc <- sweep(Sc.x, 2, SD, "/")
      vec2[grep(paste0("^", nm, "\\["), names(vec2))] <- c(Sc.x.sc)
  }

  # scale sd separately
  for(nm in SDTorescale) {
    SD.x <- vec[grep(paste0("^", nm, "\\["), names(vec))]
    vec2[grep(paste0("^", nm, "\\["), names(vec2))] <- SD.x/SD
  }
  
  # scale into sd.LV
  # sd.LV <- vec[grep("sd.LV", names(vec))]
  # vec2[grep("sd.LV", names(vec2))] <- SD*sd.LV
  
  vec2
}

RescaleChains <- function(mcmc.lst, ...) {
  rescale <- lapply(mcmc.lst, function(mcmc) {
    rot <- apply(mcmc, 1, RescaleVars, ...)
    as.mcmc(t(rot))
  })
  as.mcmc.list(rescale)
}
```


We create a new function to simulate starting values from the prior
distributions, which we can hide away

```
inits <- function(consts){
    B = matrix(rnorm(consts$nLVs*consts$NEnv),ncol=consts$nLVs)
    O = matrix(rnorm(consts$nLVs*consts$NTraits),nrow=consts$NTraits)
    varepsilon = mvtnorm::rmvnorm(consts$NSpecies,rep(0,consts$nLVs),diag(consts$nLVs))
    epsilon = mvtnorm::rmvnorm(consts$NSites,rep(0,consts$nLVs),diag(consts$nLVs))
    xi = rgamma(1,5,2)
    #for(l in 2:consts$nLVs)xi<-c(xi,rbeta(1,consts$NSpecies/(l+consts$NSpecies),l))
    xi <- c(xi,rbeta(consts$nLVs-1,1,10))
    list(
        BSTAR = B,
        OSTAR = O,
        epsilonSTAR = epsilon,
        varepsilonSTAR = varepsilon,
        sd.SiteSTAR = rexp(consts$nLVs),
        sd.SpeciesSTAR = rexp(consts$nLVs),
        beta0 = rnorm(consts$NSpecies),
        # sd.LV = rexp(consts$nLVs),
        xi = xi,
        phi = rep(1, consts$NSpecies)
    )
}
```


We use plotting functions, which are hidden here

```
PlotPost <- function(var, summ, varnames=NULL, ...) {
  var <- paste0("^", var, "*\\[")
  vars <- grep(var, rownames(summ$statistics))
  if(is.null(varnames)) varnames <- rownames(summ$statistics)[grep(var, rownames(summ$statistics))]
  if(length(varnames)!=length(vars)) 
    stop(paste0("Number of variable names, ", length(varnames), "not the same as number of variables, ",
                length(vars)))
  
  plot(summ$statistics[vars,"Mean"], 1:length(vars), 
       xlim=range(summ$quantiles[vars,]), yaxt="n", ...)
  segments(summ$quantiles[vars,"2.5%"], 1:length(vars), summ$quantiles[vars,"97.5%"], 1:length(vars))
  segments(summ$quantiles[vars,"25%"], 1:length(vars), summ$quantiles[vars,"75%"], 1:length(vars), lwd=3)
  abline(v=0, lty=3)
  axis(2, at=1:length(vars), labels=varnames, las=1)
}
AddArrows <- function(coords, marg= par("usr"), col=2) {
  origin <- c(mean(marg[1:2]), mean(marg[3:4]))
  Xlength <- sum(abs(marg[1:2]))/2
  Ylength <- sum(abs(marg[3:4]))/2
  ends <- coords / max(abs(coords)) * min(Xlength, Ylength) * .8
  arrows(
    x0 = origin[1],
    y0 = origin[2],
    x1 = ends[,
              1] + origin[1],
    y1 = ends[, 2] + origin[2],
    col = col,
    length = 0.1)

  text(
    x = origin[1] + ends[, 1] * 1.1,
    y = origin[2] + ends[, 2] * 1.1,
    labels = rownames(coords),
    col = col)
  
}
```

Latent variable models are notorious for being unidentifiable, you
can get the same mean abundances from different combinations of the
parameters. We have to make some adjustments to the model to account for
this: some of this is done in the model fitting, but for others it is
easier to do it after we obtain the posterior samples.

The details of what we do to make the HO identifiable are here

- First, we standardise \(\boldsymbol{z}\_i\) and \(\boldsymbol{\gamma}\_j\) to unit variance
  per latent variable to prevent scale invariance
- At this point the model is still invariant to sign switching:
  because \(z\_{i}\gamma\_{j} =
  (-z\_{i})(-\gamma\_{j})\) in the MCMC algorithm can (and does)
  switch signs mid run. We could solve it by placing a truncated normal
  prior on the main diagonal entries of \(\boldsymbol{B}\) or \(\boldsymbol{\omega}\), but that results in
  bimodal posterior distributions for other coefficients. Instead we
  impose sign constraints by post-process the chains. We choose one
  parameter, for example one \(z\_{i}\) or
  \(\gamma\_{j}\), to be positive, and
  switch all of the other parameters based on that one. See below for the
  details of how we fix the signs.

We want to make one parameter positive, so ideally we want to do this
to a parameter that has both modes away from zero. We can identify this
in an **ad hoc** way: for each chain for every parameter we
calculate the proportion of iterations where the sign is positive, and
then for every parameter we calculate the variance in that proportion.
If a parameter is centered around 0 the mean proportion will be about
0.5, and will not vary much, whereas if it is some way from 0 the mean
will be close to 0 or 1, and the variance will be high. ALternativley,
we coiuld look at the largest \(|p -
0.5|\), or we can vissually identify such parameters from the
posterior samples. There may be even better alternatives.

#### Maximum informed dimensions

Since we have two or more latent variables, we also have to worry about
their rotation. In the previous example we fixed the rotation by setting
some parameters to zero, but here we do this differently.

For those who are interested, this is explained further here.

Previously, we set some parameters to zero in the environment effects
\(\symbf{B}\) and trait effects \(\symbf{\omega}\) to fix the rotation.
Fixing parameters has as advantage that the model is clearly
(mathematically) identifiable. However, it has the downside of
introducing order dependence in the predictors, traits, and species
effects. Bhattacharya
and Dunson (2011) developed a multiplicative gamma shrinkage prior
which prevents this order dependence, though it leads less clearly to a
mathematical identifiability so that (for ordination) the results need
to be post-processed so that all MCMC iterations have the same rotation.
In this vignette we use a similar prior as Bhattacharya
and Dunson (2011), though based on a product of a Gamma and of beta
distributions. Use of a beta priors introduces an order constraint to
the variance of the latent variables, so that it can only decrease, and
has the intuitive interpretation of proportional decrease in variance of
one latent variable to the next.

The infinite factor model is motivated by the idea that, although the
latent variables may be affected by the rotation, the product of species
loadings (i.e., the species associations) is not. The main effects for
the predictors, as well as the fourth corner term, are invariant to the
rotation and can be estimated even when the model is unidentifiable. The
choice of the number of latent variables may be more important in order
for an accurate representation of patterns.

First we introduce the additional \(d-\)sized vector \(\symbf{\xi}\), with \(\xi\_1 \sim \Gamma(5, 2)\) and \(\xi\_{q \in d \setminus \{1\}} \sim
B(p/(2q+p),q)\) (or alternatively, to introduce less shrinkage
e.g., \(B(5, 5)\) or even \(U(0,1)\)). We use this vector to construct
the LV variation parameters as: \(\Sigma\_{q,q}
= \prod^d\_{q=1} \delta\_{1:q}\), so that each latent variable
explains less variation than the next, as long as the \(\Sigma\_{q,q}\) are distinctly different.
The choice of gamma distribution is purposefully uninformative, since we
do not know how much variation the latent variables explain in the data
(but usually, this ranges between 1-5). The choice of parameterization
for the beta distribution is motivated as a shrinkage prior; with a low
mean and large variance so that the variation for each consecutive
latent variable is forced to be considerably lower than that of the
previous latent variable. Additionally, by using the index of the latent
variable, and the total number of species, as parameters for the prior
we ensure that each latent variable receives more shrinkage than the
next, and emphasis is placed on a relatively small number of latent
variables.

We could still estimate *some* but not all latent variables,
for example in larger examples and to speed up computation. The maximum
number of latent variables that can be informed (see van der Veen et
al. (2023)) by the predictor variables is determined by the number of
traits and environmental predictors that we have. In essence, we cannot
have more latent variables that the minimum number of traits and
environmental predictors, because at that point we have reached the
maximum amount of information that (one of, or the combination of) the
matrices can explain in the responses (i.e., one of the matrices alone
could explain more variation still). In the example here, there are five
environmental predictors, but seven traits. Thus, the maximum number of
latent variables that can be informed by the matrices jointly is five,
though two more dimensions could be informed by the traits alone.

The model code is included here

```
# Model code
# Update our constants from before with the new number of LVs, rest remains the same
HO.nim <- nimbleCode({
  for (i in 1:NSites) {
    for (j in 1:NSpecies) {
      Y[i, j] ~ dnegbin(phi[j]^-1/(phi[j]^-1+lambda[i, j]), phi[j]^-1)
    }      
  }

    ones[1:NSites] <- rep(1, NSites)
    # linear predictor
    log(lambda[1:NSites, 1:NSpecies]) <- eta[1:NSites, 1:NSpecies]
    eta[1:NSites,1:NSpecies] <-  asCol(ones[1:NSites])%*%asRow(beta0[1:NSpecies]) +z[1:NSites,1:nLVs]%*%Sigma[1:nLVs,1:nLVs]%*%t(gamma[1:NSpecies,1:nLVs])
    Sigma[1:nLVs, 1:nLVs] <- diag(sd.LV[1:nLVs])
    # sites
    XB[1:NSites,1:nLVs] <- X[1:NSites,1:NEnv]%*%B.ort[1:NEnv,1:nLVs]
    zSTAR[1:NSites,1:nLVs] <- XB[1:NSites,1:nLVs] + epsilonSTAR[1:NSites,1:nLVs]%*%diag(sd.SiteSTAR[1:nLVs])
    # species
    gammaSTAR[1:NSpecies,1:nLVs] <- omegaTR[1:NSpecies,1:nLVs] + varepsilonSTAR[1:NSpecies,1:nLVs]%*%diag(sd.SpeciesSTAR[1:nLVs])
    omegaTR[1:NSpecies,1:nLVs] <- TR[1:NSpecies,1:NTraits]%*%O.ort[1:NTraits,1:nLVs]
  
  # prior for intercept
  beta0[1:NSpecies] ~ dmnorm(zeroNSpecies[1:NSpecies], prec = diagNSpecies[1:NSpecies,1:NSpecies])
  
  # prior for LV scales
  xi[1] ~ dgamma(5,2)
  for(l in 2:nLVs) {
  xi[l] ~ dbeta(1, 10)#NSpecies/(2*l+NSpecies), l^2) #dbeta(1,20)# could instead be dunif(0,1) to be less informative
  }
  
  # diagonal matrices and such for use in priors
  diagNTraits[1:NTraits,1:NTraits] <- diag(NTraits)
  diagNEnv[1:NEnv,1:NEnv] <- diag(NEnv)
  zeroNTraits[1:NTraits] <- rep(0, NTraits)
  zeroNEnv[1:NEnv] <- rep(0, NEnv)
  diagNSites[1:NSites,1:NSites] <- diag(NSites)
  diagNSpecies[1:NSpecies,1:NSpecies] <- diag(NSpecies)
  zeroNSites[1:NSites] <- rep(0, NSites)
  zeroNSpecies[1:NSpecies] <- rep(0, NSpecies)
  
    for(l in 1:nLVs) { 
    # prors for LV-level errors
    epsilonSTAR[1:NSites,l] ~ dmnorm(zeroNSites[1:NSites], prec = diagNSites[1:NSites,1:NSites])# Residual
    varepsilonSTAR[1:NSpecies,l] ~ dmnorm(zeroNSpecies[1:NSpecies], prec = diagNSpecies[1:NSpecies,1:NSpecies]) # Residual
    # priors for predictor effects
    OSTAR[1:NTraits,l] ~ dmnorm(zeroNTraits[1:NTraits], prec = diagNTraits[1:NTraits,1:NTraits]) 
    BSTAR[1:NEnv,l] ~ dmnorm(zeroNEnv[1:NEnv], prec = diagNEnv[1:NEnv,1:NEnv]) 
    # priors for scale parameters
    sd.SiteSTAR[l] ~ dexp(1)
    sd.SpeciesSTAR[l] ~ dexp(1)
    # constructing LV scale
    sd.LV[l] <- prod(xi[1:l])
        
    ## standardizing z and gamma for identifiability
    ## and B and O to have non-unit norms
    ## rescale  the rest of the parameters for trace plots and such
    ## these are the quantities that really go into eta
    #norm.B[l] <- sqrt(sum(BSTAR[1:NEnv,l]^2))
    #norm.O[l] <- sqrt(sum(OSTAR[1:NTraits,l]^2))
    StdDev.z[l] <- sd(zSTAR[1:NSites,l])
    z[1:NSites,l] <- zSTAR[1:NSites,l]/StdDev.z[l]
    epsilon[1:NSites,l] <- epsilonSTAR[1:NSites,l]/StdDev.z[l]
    B[1:NEnv,l] <- B.ort[1:NEnv,l]/StdDev.z[l]#*norm.B[l]
    sd.Site[l] <- sd.SiteSTAR[l]/StdDev.z[l]
    
    StdDev.gamma[l] <- sd(gammaSTAR[1:NSpecies,l])
    gamma[1:NSpecies,l] <- gammaSTAR[1:NSpecies,l]/StdDev.gamma[l]
    varepsilon[1:NSpecies,l] <- varepsilonSTAR[1:NSpecies,l]/StdDev.gamma[l]
    O[1:NTraits,l] <- O.ort[1:NTraits,l]/StdDev.gamma[l]#*norm.O[l]
    sd.Species[l] <- sd.SpeciesSTAR[l]/StdDev.gamma[l]
    }
  
    # Orthogonalize B and O for identifiability
    # first predictor effects B by its F norm
    # B.nn[1:NEnv,1:nLVs] <- BSTAR[1:NEnv,1:nLVs]%*%diag(1/norm.B[1:nLVs])
    # O.nn[1:NTraits,1:nLVs] <- OSTAR[1:NTraits,1:nLVs]%*%diag(1/norm.O[1:nLVs])
    # # get Cholesky factor
    # L.B[1:nLVs,1:nLVs] <- chol(t(B.nn[1:NEnv,1:nLVs])%*%B.nn[1:NEnv,1:nLVs])
    # L.O[1:nLVs,1:nLVs] <- chol(t(O.nn[1:NTraits,1:nLVs])%*%O.nn[1:NTraits,1:nLVs])
    # # Orthogonalize predictor effects
    # B.ort[1:NEnv,1:nLVs] <- t(forwardsolve(t(L.B[1:nLVs,1:nLVs]),t(B.nn[1:NEnv,1:nLVs])))
    # O.ort[1:NTraits,1:nLVs] <- t(forwardsolve(t(L.O[1:nLVs,1:nLVs]),t(O.nn[1:NTraits,1:nLVs])))
    B.ort[1:NEnv,1:nLVs] <- nimQR.u(BSTAR[1:NEnv,1:nLVs], nLVs)
    O.ort[1:NTraits,1:nLVs] <- nimQR.u(OSTAR[1:NTraits,1:nLVs],nLVs)
        for (j in 1:NSpecies) {
     phi[j] ~ dexp(1)
    }
})
# QR schwarz-Rutishauser
# The algorithm is correct, but I cannot use it with HMC..
nimQR.u <- nimbleFunction(
  run = function(A = double(2), p  = double()) {
    for (k in 2:p) {
      A[, k] <- A[, k]
      for (i in 1:(k-1)) {
        A[, k] <- A[, k] - inprod(A[,i], A[,k]) / inprod(A[, i], A[, i]) * A[, i]
      }
    }
    return(A)
    returnType(double(2))
  } ,
buildDerivs = list(run = list(ignore = c("k", "i","p")))
)
```

And now we can run the model:

```
consts$nLVs <- nLVs <- 10
HO.LV <- ParaNimble(4, fun = RunOneChain,
                       dat = dat,
                       code = HO.nim,
                       inits = inits, 
#                       Nburn=5e1, NIter=5.5e2, Nthin=1, # for a small run
                       Nburn = 1500, NIter = 15e3, Nthin = 1, # for a full run
                       consts = consts, block = MakeBlockList(consts, LVwise = FALSE), slice = c(paste0("beta0[",1:consts$NSpecies,"]"),paste0("phi[",1:consts$NSpecies,"]"))) # HMC on the rest (hypers)
```

The MCMC takes a Looooonnnggg time to run (= 2 weeks), so we have
saved the output here.
You will need to download it into the same file as the Rmd file if you
want to run this (we have tried to write code to do this, and yes that
includes trying dl=1 and raw=1, but that doesn’t work).

```
consts$nLVs <- nLVs <- 10 # because the last chunk is not evaluated
VarsToProcess <- c("^BSTAR", "^OSTAR", "B", "O", "^epsilonSTAR", "^varepsilonSTAR")

# post-process chains for sign-swapping
if(exists("HO.LV")) {
  chains2LV <- postProcess(HO.LV, VarsToProcess = VarsToProcess, rule = 2, print = TRUE)
} else {
  load("chains2LV.RData")
}

# Calculate summaries
summ.HOLV <- summary(chains2LV)
```

#### Results

First, we look at the LV variation. After all, that is what is different
in this vignette.

```
library(basicMCMCplots)
chainsPlot(chains2LV, var = c("sd.LV"), legend = F, traceplot=TRUE)
```

We can also plot these as a sort of “screeplot”, to see how the
variation of the LVs decreases with the number of dimensions:

```
par(mfrow=c(1,2))
plot(NA,ylim=c(0, max(rbind(summ.HOLV$statistics[grep("sd.LV", row.names(summ.HOLV$statistics)),1],
                            summ.HOLV$quantiles[grep("sd.LV", row.names(summ.HOLV$statistics)),1], 
                            summ.HOLV$statistics[grep("sd.LV", row.names(summ.HOLV$statistics)),4]))+0.1),
     xlim=c(1,consts$nLVs), xlab="Latent variable", ylab="SD")
lines(summ.HOLV$statistics[grep("sd.LV", row.names(summ.HOLV$statistics)),1], 
      type="b",xlab="Latent variable",ylab="SD")
lines(summ.HOLV$quantiles[grep("sd.LV", row.names(summ.HOLV$statistics)),1], 
      type="b",xlab="Latent variable",ylab="SD",col="red",lty="dashed")
lines(summ.HOLV$quantiles[grep("sd.LV", row.names(summ.HOLV$statistics)),4], 
      type="b",xlab="Latent variable",ylab="SD",col="red",lty="dashed")

plot(NA,ylim=c(0,max(rbind(summ.HOLV$statistics[grep("xi", row.names(summ.HOLV$statistics)),1], 
                           summ.HOLV$quantiles[grep("xi", row.names(summ.HOLV$statistics)),1], 
                           summ.HOLV$statistics[grep("xi", row.names(summ.HOLV$statistics)),4]))+0.1), 
     xlim=c(1,consts$nLVs),xlab="Latent variable",ylab="xi")
lines(summ.HOLV$statistics[grep("xi", row.names(summ.HOLV$statistics)),1], 
      type="b",xlab="Latent variable",ylab="xi")
lines(summ.HOLV$quantiles[grep("xi", row.names(summ.HOLV$statistics)),1],
      type="b",xlab="Latent variable",ylab="xi",col="red",lty="dashed")
lines(summ.HOLV$quantiles[grep("xi", row.names(summ.HOLV$statistics)),4],
      type="b",xlab="Latent variable",ylab="xi",col="red",lty="dashed")
```


Next, we look at the mixing of the predictor coefficients.

Environment first:

```
library(basicMCMCplots)
chainsPlot(chains2LV, var = c("B"), legend = F, traceplot=TRUE)
```

And the trait effects:

```
chainsPlot(chains2LV, var = c("O"), legend = F, traceplot=TRUE)
```

To get a better impression of the uncertainty of the effects, we also
look at them in a caterpillar plot:

```
par(mar=c(5, 11, 4, 2) + 0.1)
PlotPost("B",summ.HOLV,varnames = paste(rep(colnames(X),consts$nLVs), rep(paste0("LV.",1:consts$nLVs),each=ncol(X)), sep = "-"), xlab="Predictor effect", ylab=NA)
```

```
par(mar=c(5, 11, 4, 2) + 0.1)
PlotPost("O",summ.HOLV,varnames = paste(rep(colnames(TR),consts$nLVs), rep(paste0("LV.",1:consts$nLVs),each=ncol(TR)), sep = "-"), xlab="Predictor effect", ylab=NA)
```


What we are really interested in, is looking at the environment-trait
interactions. We can plot those with the following code

.

First we get the interactions:

```
# Function to get MCMC results for reduced-rank approximated interaction term
getMCMCInt <- function(mcmc.lst, dat,...) {
  int.mcmc <- lapply(mcmc.lst, function(mcmc){
    inCoefs <- as.mcmc(t(apply(mcmc, 1, function(ch){
    B <- ChainToMatrix(ch,"B")
    O <- ChainToMatrix(ch,"O")  
    SDs <- diag(ch[grep("sd.LV",gsub("\\s*\\[[^\\)]+\\]","",names(ch)))])
    coefs <- B%*%SDs%*%t(O)
    vec <- c(coefs)
    names(vec) <- paste0(colnames(dat$X),":",rep(colnames(dat$TR),each=ncol(dat$X)))
    vec
    })))
  }
  )
  as.mcmc.list(int.mcmc)
}

intChains <- getMCMCInt(chains2LV, dat = dat)
```

Second, we plot them.

```
summ.EnvTraitInt <- summary(intChains)
coefs <- matrix(summ.EnvTraitInt$statistics[,1], ncol = ncol(dat$TR), nrow=ncol(dat$X), dimnames = list(colnames(dat$X),colnames(dat$TR)))
a <- 0.2
colort <- colorRampPalette(c("blue", "white", "red"))
lattice::levelplot(coefs, xlab = "Environmental Variables", 
                      ylab = "Species traits", col.regions = colort(100), cex.lab = 1.3, 
                      at = seq(-a, a, length = 100), scales = list(x = list(rot = 45)))
```

Now, we can make two-dimensional ordination plots of sites and
species, with their predictor effects. We scale both the site scores and
species loadings by the square root of the LV variation.

```
# get LV variation for plotting
SDs <- summ.HOLV$statistics[grep("sd.LV", rownames(summ.HOLV$statistic)),]
rownames(SDs) <- c(paste0("Latent Variable ", 1:10))

sd.SiteSTAR <- GetMeans(summ.HOLV, name="^sd.SiteSTAR", d=consts$nLVs)
sd.SpeciesSTAR <- GetMeans(summ.HOLV, name="^sd.SpeciesSTAR", d=consts$nLVs)

varepsilon.STAR <- GetMeans(summ.HOLV, name="^varepsilonSTAR", d=consts$nLVs)%*%diag(c(sd.SpeciesSTAR))
epsilon.STAR <- GetMeans(summ.HOLV, name="^epsilonSTAR", d=consts$nLVs)%*%diag(c(sd.SiteSTAR))
BSTAR <- GetMeans(summ.HOLV, name="BSTAR", d=consts$nLVs)
row.names(BSTAR) <- colnames(dat$X)
OSTAR<- GetMeans(summ.HOLV, name="OSTAR", d=consts$nLVs)
row.names(OSTAR) <- colnames(dat$TR)

# Need to orthogonalize the means for BSTAR and OSTAR
B.ort <- nimQR.u(BSTAR, consts$nLVs)
O.ort <- nimQR.u(OSTAR, consts$nLVs)

# Form site scores and loadings, standardize them
# Scaling, same as ^alpha=0.5 in a biplot, equal weight for sites and species
zSTAR <- X%*%B.ort+epsilon.STAR
z.sd <- apply(zSTAR,2,sd)
z.m <- zSTAR%*%solve(diag(z.sd))%*%diag(sqrt(SDs[,1]))
B.m <- B.ort%*%solve(diag(z.sd))%*%diag(sqrt(SDs[,1]))
  
gammaSTAR <- TR%*%O.ort + varepsilon.STAR
gamma.sd <- apply(gammaSTAR,2,sd)
gamma.m <- gammaSTAR%*%solve(diag(gamma.sd))
O.m <- O.ort%*%solve(diag(gamma.sd))%*%diag(sqrt(SDs[,1]))

# variance explained
1-sum(apply(varepsilon.STAR%*%solve(diag(gamma.sd))%*%diag(sqrt(SDs[,1])),2,var))/sum(SDs[,1]^2) #47% explained by traits
```

```
## [1] 0.4700789
```

```
1-sum(apply(epsilon.STAR%*%solve(diag(z.sd))%*%diag(sqrt(SDs[,1])),2,var))/sum(SDs[,1]^2) #75 by environment
```

```
## [1] 0.7570908
```

```
apply(epsilon.STAR%*%solve(diag(z.sd))%*%diag(sqrt(SDs[,1])),2,var)/sum(SDs[,1]^2)
```

```
##  [1] 4.195825e-03 4.510831e-02 8.485021e-02 3.970619e-02 3.459327e-02
##  [6] 1.224336e-02 1.085495e-02 1.084027e-02 4.845128e-04 3.229258e-05
```

```
apply(epsilon.STAR%*%solve(diag(z.sd)),2,var)/sum(apply(epsilon.STAR%*%solve(diag(z.sd)),2,var))
```

```
##  [1] 0.007746183 0.111644249 0.260652604 0.144750491 0.147668960 0.062717793
##  [7] 0.071602658 0.127138477 0.038467345 0.027611241
```

```
apply(varepsilon.STAR%*%solve(diag(gamma.sd)),2,var)/sum(apply(varepsilon.STAR%*%solve(diag(gamma.sd)),2,var))
```

```
##  [1] 0.11913632 0.12880975 0.14320562 0.16780726 0.20580285 0.09073037
##  [7] 0.05304759 0.06346019 0.01266010 0.01533995
```

```
#LVs
plot(z.m[,1:2],type="n", xlab="Latent variable 1", ylab="Latent variable 2", main = "Sites")
text(z.m[,1:2],labels=1:consts$NSites)
AddArrows(marg = par("usr"), coords=B.m[,1:2], col="red")

#gammas
plot(gamma.m[,1:2],type="n", xlab="Latent variable 1", ylab="Latent variable 2", main = "Species")
text(gamma.m[,1:2],labels=vegan::make.cepnames(colnames(Y)))
AddArrows(marg = par("usr"), coords=O.m[,1:2], col="blue")
```

These tell us how well the predictors explain the ordination; the
species- and site-specific scale parameters would be zero if the
predictors fully explained the ordination. The scale parameters for the
latent variables are similar to singular values (the square root of
eigenvalues) in a classical ordination; they reflect a dimension its
importance to the response.

#### Residual covariance matrix


The maths behind this is covered in here


\[\begin{multline}
\text{cov}(\boldsymbol{z}\_i^\top\boldsymbol{\Sigma}\boldsymbol{\gamma}\_j,
\boldsymbol{z}\_{i2}^\top\boldsymbol{\Sigma}\boldsymbol{\gamma}\_{j2}) =
\text{cov}(\boldsymbol{x}\_i^\top\boldsymbol{B}\boldsymbol{\Sigma}\boldsymbol{\varepsilon}\_j,\boldsymbol{x}\_{i2}^\top\boldsymbol{B}\boldsymbol{\Sigma}\boldsymbol{\varepsilon}\_{j2})
+
\text{cov}(\boldsymbol{x}\_i^\top\boldsymbol{B}\boldsymbol{\Sigma}\boldsymbol{\varepsilon}\_j,\boldsymbol{tr}\_{j2}^\top\boldsymbol{\omega}\boldsymbol{\Sigma}\boldsymbol{\epsilon}\_{i2})
+
\text{cov}(\boldsymbol{x}\_i^\top\boldsymbol{B}\boldsymbol{\Sigma}\boldsymbol{\varepsilon}\_j,\boldsymbol{\epsilon}\_{i2}^\top\boldsymbol{\Sigma}\boldsymbol{\varepsilon}\_{j2})
+
\text{cov}(\boldsymbol{tr}\_j^\top\boldsymbol{\omega}\boldsymbol{\Sigma}\boldsymbol{\epsilon}\_i,\boldsymbol{x}\_{i2}^\top\boldsymbol{B}\boldsymbol{\Sigma}\boldsymbol{\varepsilon}\_{j2})
+
\text{cov}(\boldsymbol{tr}\_{j}^\top\boldsymbol{\omega}\boldsymbol{\Sigma}\boldsymbol{\epsilon}\_{i},\boldsymbol{tr}\_{j2}^\top\boldsymbol{\omega}\boldsymbol{\Sigma}\boldsymbol{\epsilon}\_{i2})
+ \\
\text{cov}(\boldsymbol{tr}\_j^\top\boldsymbol{\omega}\boldsymbol{\Sigma}\boldsymbol{\epsilon}\_i,\boldsymbol{\epsilon}\_{i2}^\top
\boldsymbol{\Sigma}\boldsymbol{\varepsilon}\_{j2}) +
\text{cov}(\boldsymbol{\epsilon}\_{i}^\top
\boldsymbol{\Sigma}\boldsymbol{\varepsilon}\_{j},\boldsymbol{x}\_{i2}^\top\boldsymbol{B}\boldsymbol{\Sigma}\boldsymbol{\varepsilon}\_{j2})
+
\text{cov}(\boldsymbol{\epsilon}\_{i}^\top \boldsymbol{\Sigma}
\boldsymbol{\varepsilon}\_{j},\boldsymbol{tr}\_{j2}^\top\boldsymbol{\omega}\boldsymbol{\Sigma}\boldsymbol{\epsilon}\_{i2})+
\text{cov}(\boldsymbol{\epsilon}\_i^\top
\boldsymbol{\Sigma}\boldsymbol{\varepsilon}\_j,\boldsymbol{\epsilon}\_{i2}^\top
\boldsymbol{\Sigma}\boldsymbol{\varepsilon}\_{j2}),
\end{multline}\] third order terms are zero for central normal
random variables, so this simplifies to: \[\begin{equation}
\text{cov}(\boldsymbol{z}\_i^\top\boldsymbol{\Sigma}\boldsymbol{\gamma}\_j,
\boldsymbol{z}\_{i2}^\top\boldsymbol{\Sigma}\boldsymbol{\gamma}\_{j2}) =
\boldsymbol{x}\_i^\top\boldsymbol{B}\boldsymbol{\Sigma}\text{cov}(\boldsymbol{\varepsilon}\_j,\boldsymbol{\varepsilon}\_{j2})
\boldsymbol{\Sigma}\boldsymbol{B}^\top\boldsymbol{x}\_{i2}+
\boldsymbol{x}\_i^\top\boldsymbol{B}\boldsymbol{\Sigma}\text{cov}(\boldsymbol{\varepsilon}\_j,\boldsymbol{\epsilon}\_{i2})\boldsymbol{\Sigma}\boldsymbol{\omega}^\top\boldsymbol{tr}\_{j2}
+
\boldsymbol{tr}\_j^\top\boldsymbol{\omega}\boldsymbol{\Sigma}\text{cov}(\boldsymbol{\epsilon}\_i,\boldsymbol{\varepsilon}\_{j2})\boldsymbol{\Sigma}\boldsymbol{B}^\top\boldsymbol{x}\_{i2}
+
\boldsymbol{tr}\_{j}^\top\boldsymbol{\omega}\boldsymbol{\Sigma}\text{cov}(\boldsymbol{\epsilon}\_{i},\boldsymbol{\epsilon}\_{i2})\boldsymbol{\Sigma}\boldsymbol{\omega}^\top\boldsymbol{tr}\_{j2}
+
\text{cov}(\boldsymbol{\epsilon}\_i^\top
\boldsymbol{\Sigma}\boldsymbol{\varepsilon}\_j,\boldsymbol{\epsilon}\_{i2}^\top
\boldsymbol{\Sigma} \boldsymbol{\varepsilon}\_{j2})
\end{equation}\]

Due to independence of all species and site effects, \(\text{cov}(\boldsymbol{\varepsilon}\_j,\boldsymbol{\epsilon}\_{i2})
= \text{cov}(\boldsymbol{\epsilon}\_i,\boldsymbol{\varepsilon}\_{j2}) =
0\), \(\text{cov}(\boldsymbol{\epsilon}\_i,
\boldsymbol{\epsilon}\_{i2}) = 0\), and \(\text{cov}(\boldsymbol{\varepsilon}\_j,
\boldsymbol{\varepsilon}\_{j2}) = 0\), so that the residual
covariance matrix for the full residual vector of all sites and species
is:

\[\begin{equation}
\text{cov}\{\text{vec}(\boldsymbol{Z}\boldsymbol{\Sigma}\boldsymbol{\Gamma}^\top),
\text{vec}(\boldsymbol{Z}\boldsymbol{\Sigma}\boldsymbol{\Gamma}^\top)\}
=
\textbf{TR}\boldsymbol{\omega}\boldsymbol{\Sigma}\text{diag}(\boldsymbol{\delta}^2)\boldsymbol{\Sigma}\boldsymbol{\omega}^\top\textbf{TR}^\top
\otimes
\textbf{X}\textbf{B}\boldsymbol{\Sigma}\text{diag}(\boldsymbol{\delta}^2)\boldsymbol{\Sigma}\textbf{B}^\top\textbf{X}^\top.
\end{equation}\]

Here, the residual covariances are visualized for all species on the
first site:

```
Sigma <- diag(SDs[grep("Latent Variable", rownames(SDs)),1])
Sigma.sp <- diag(c(sd.SpeciesSTAR/gamma.sd))^2
Sigma.si <- diag(c(sd.SiteSTAR/z.sd))^2

covMat <- TR%*%O.m%*%Sigma%*%Sigma.si%*%Sigma%*%t(O.m)%*%t(TR)*(X%*%B.m%*%Sigma%*%Sigma.sp%*%t(B.m)%*%t(X))[1]
colnames(covMat) <- row.names(covMat) <- colnames(Y)
par(mfrow=c(1,1))
corrplot::corrplot(cov2cor(covMat),type = "lower",order = "AOE", main = "Species", mar = c(1,1,1,1),tl.srt=45,tl.cex = .5)
```
